# Onsite fixation preserves phosphoprotein profile in human whole blood

**DOI:** 10.64898/2026.11.05.763716

**Authors:** Lily Shang, McKenna Coughlin, Carly Lancaster, Angelynn Huang, Justin A. Jarrell, Li-Chun Cheng

## Abstract

Phospho-flow cytometry is increasingly used as a pharmacodynamic readout of kinase-pathway activity, including in JAK inhibitor trials. Phosphoprotein states are intrinsically labile *ex vivo* and are further altered by conventional workflows, which typically involve PBMC isolation, cryopreservation, and transit times of hours to days before measurement. Here, we fix whole blood at collection and quantify six phosphoproteins by mass cytometry, benchmarked against matched cryopreserved PBMCs. Unlike PBMCs, whole blood retained neutrophils, ∼75–80% of which were pSTAT3-positive at baseline. Basal pSTAT3 was also markedly higher in whole blood monocytes (∼41% positive vs. ∼9% in PBMCs), whereas T, B, and NK cells differed by less than 1 percentage point. Stimulation produced subset-specific phosphorylation patterns that were retained after 6 weeks of storage. Because fixation does not require centrifugation and retains neutrophils, samples can be fixed at collection sites without on-site processing laboratories and analyzed centrally in batches.

**Highlights:** Fixing whole blood at the draw site preserves basal phosphoprotein states Six phosphoproteins measured by mass cytometry in subsets including neutrophils Basal pSTAT3+ monocytes: ∼41% in fixed whole blood versus ∼9% in PBMCs Phospho-epitopes in fixed whole blood remain stable for 6 weeks of storage

**Graphical abstract:** 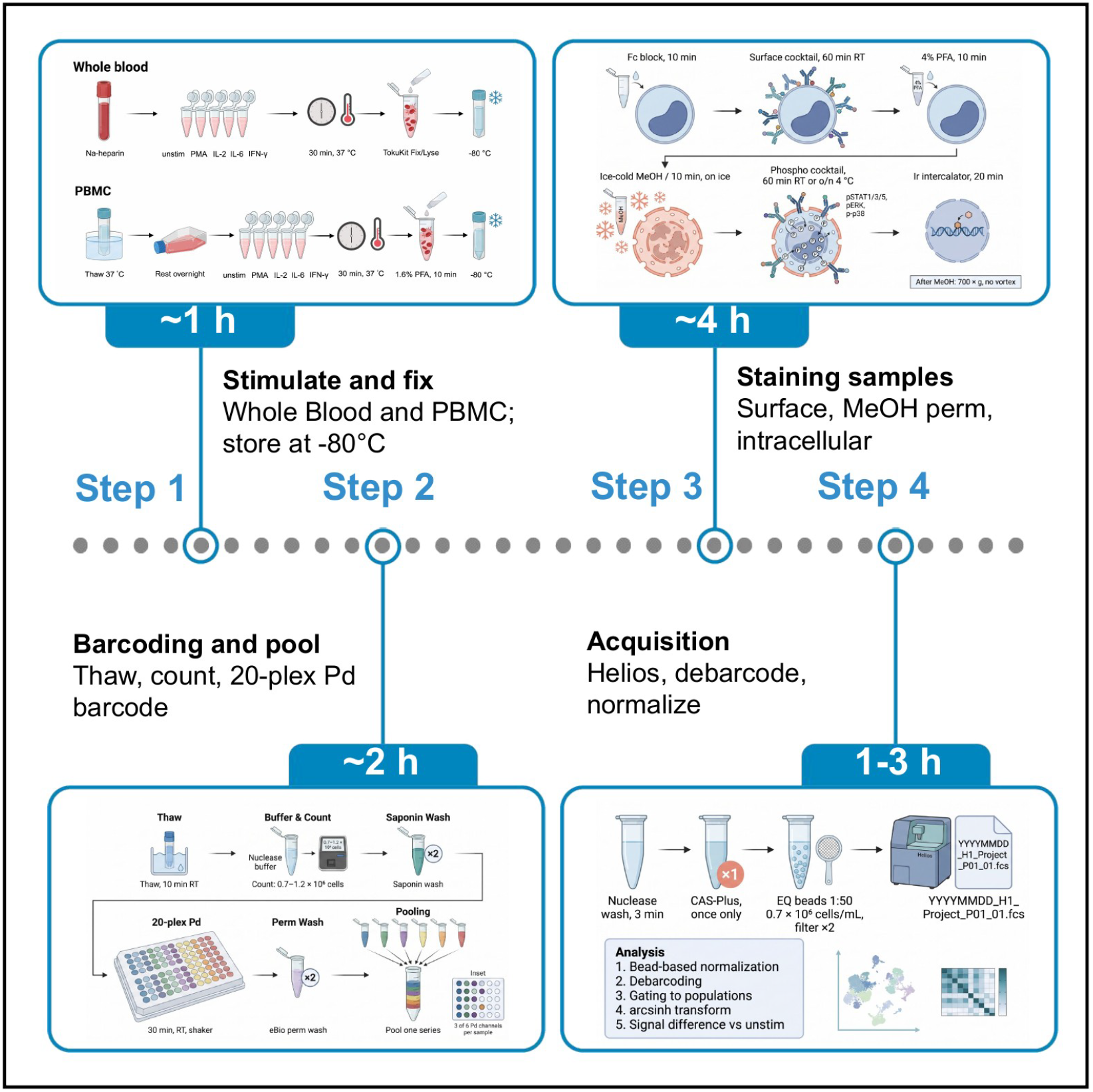

## Introduction

Phosphorylation-based readouts serve as key pharmacodynamic endpoints in early-phase clinical trials, reporting target engagement and downstream pathway modulation directly in patient samples. Annual JAK inhibitor trial counts increased from 61 in 2015 to 206 in 2024.^1^ Because phosphoprotein epitopes are dynamic and susceptible to preanalytical variability, rapid cell fixation is essential to prevent enzymatic dephosphorylation or stress-induced signaling cascades.^2^ Subsequent methanol-based permeabilization gives antibodies access to nuclear and cytoplasmic epitopes. Because fixatives and permeabilizing agents can alter binding sites, each panel requires optimization and validation.^2^

Current workflows favor peripheral blood mononuclear cell (PBMC) isolation and cryopreservation, which allows on-site biobanking and batch processing in a central laboratory. However, ambient handling, centrifugation, and mechanical shear stress during PBMC isolation induce artifactual signaling and alter basal phosphoprotein levels before fixation.^3^ Cryopreservation and freeze-thaw cycles further deplete granulocytes and shift immune subset frequencies.^4^ In multi-site clinical trials, shipping unpreserved specimens delays processing. Basal signals then degrade to the point that *ex vivo* stimulation is often required to detect a phospho-signal.^3,4^ PBMC isolation also removes neutrophils, along with any phospho-signaling they carry.

Here, we fix whole blood with a paraformaldehyde (PFA)-based stabilization kit (TokuKit) immediately after the draw. Immediate fixation captures basal phosphoprotein levels without *ex vivo* stimulation, and an optional short stimulation can precede fixation. Fixed samples are stored at −80°C until batched analysis, which avoids granulocyte loss, centrifugation stress, and freeze-thaw of viable cells. We quantified six phosphoproteins across major peripheral immune subsets in stabilized whole blood and in matched cryopreserved PBMCs. The targets were p38 (T180/Y182), pERK1/2 (T202/Y204), pS6 (S235/S236), pSTAT1 (Y701), pSTAT3 (Y705), and pSTAT5 (Y694).

Compared with PBMC workflows, this protocol changes three things. First, fixation and red blood cell lysis happen in one kit at the collection site, with no centrifugation before fixation. Second, neutrophils, which PBMC isolation removes, remain available for phospho-analysis. Third, the same validated panel runs on fixed whole blood and cryopreserved PBMCs, so new whole blood data can be compared directly with existing PBMC datasets.

## Results

### Paired whole blood and PBMC phospho-profiling workflow

We tested five signaling conditions per sample type: basal control, PMA/ionomycin, IL-2, IL-6, and IFN-γ (Table S1). Whole blood was fixed in TokuKit with concurrent red blood cell lysis, and PBMCs were fixed in 1.6% PFA (Figure 1A). After storage at −80°C, samples were barcoded with a 20-plex palladium set, surface stained, permeabilized with methanol, and stained for phospho-epitopes (Figure 1B).

**Figure 1.**
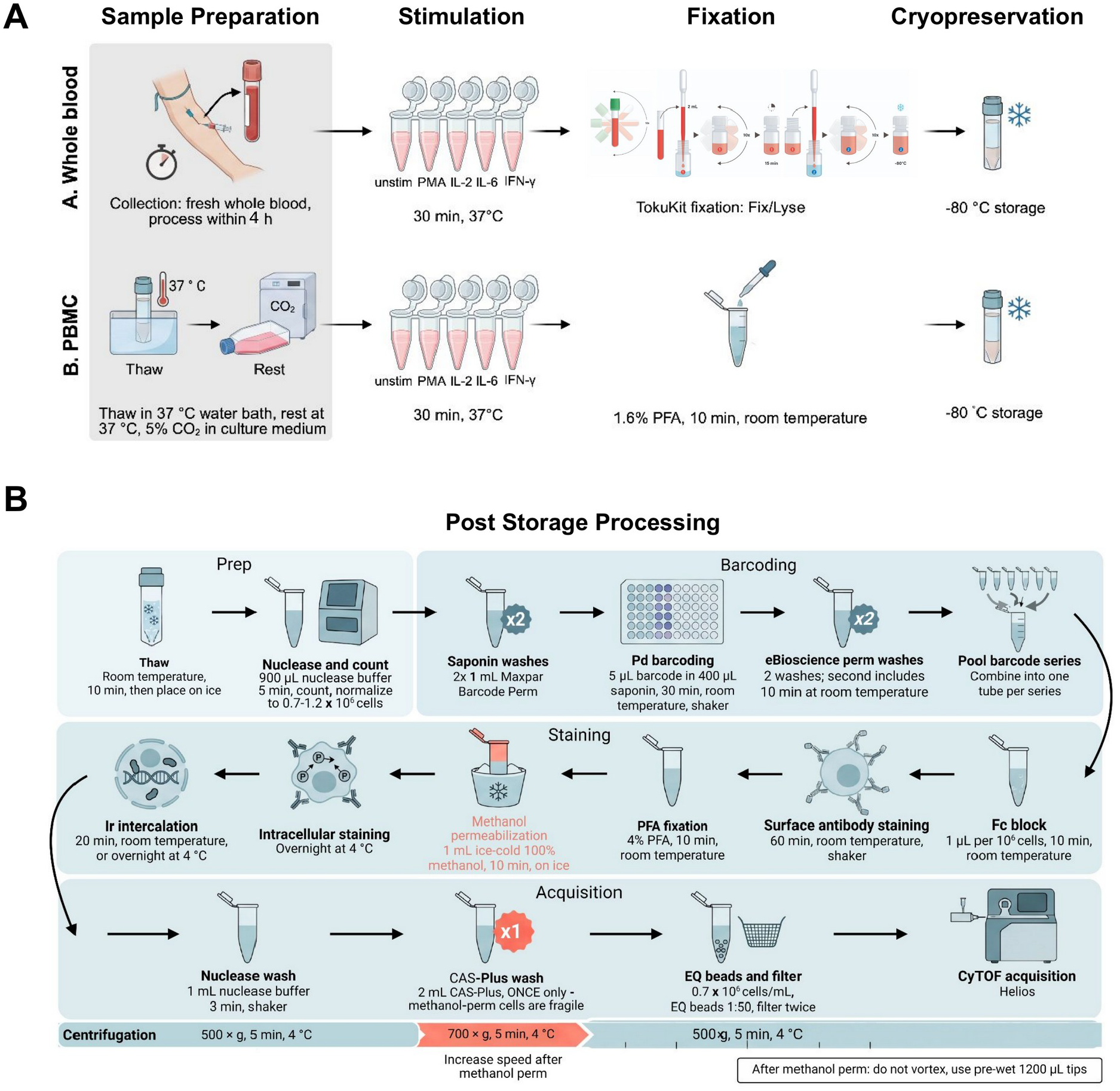
Sample processing workflow for whole blood and PBMCs. **(A)** Parallel processing of whole blood and PBMC. Both sample types are stimulated for 30 min at 37°C with PMA/ionomycin, IL-2, IL-6, IFN-γ, or left unstimulated. Whole blood samples are fixed in TokuKit with concurrent red blood cell lysis. PBMCs are fixed in 1.6% PFA for 10 min at room temperature. Fixed samples are cryopreserved at −80°C prior to barcoding and staining (B). **(B)** Post-storage workflow from thaw through CyTOF acquisition, including palladium barcoding, surface and intracellular staining, methanol permeabilization, and iridium intercalation. Centrifugation speed is increased to 700 × g after methanol permeabilization because permeabilized cells are fragile. The post-permeabilization CAS-Plus wash is limited to one wash to minimize cell loss.

Data were acquired on a Helios mass cytometer with the antibody panel in Table S2. Antibody clones validated in fixed whole blood are listed at teiko-labs.com/tokukit.

Gating resolved 10 cell populations from fixed whole blood: neutrophils, basophils and plasmacytoid Dendritic Cells (pDCs), T cells, B cells, cytokine-producing and cytolytic natural killer (NK) cells, classical, intermediate, and non-classical monocytes, and conventional DCs (cDCs) (Figure S1).

### Stimulation induces subset-specific phosphorylation in fixed whole blood

In whole blood leukocytes, PMA/ionomycin increased p38-positive cells from 28.31% to 80.88%, pERK1/2-positive cells from 1.77% to 88.79%, and pS6-positive cells from 1.94% to 62.19% (Figure 2A). IL-6 increased pSTAT3-positive cells from 55.75% to 81.36% and pSTAT1-positive cells from 12.72% to 32.75%. IL-2 increased pSTAT5-positive cells from 1.45% to 9.52%.

**Figure 2.**
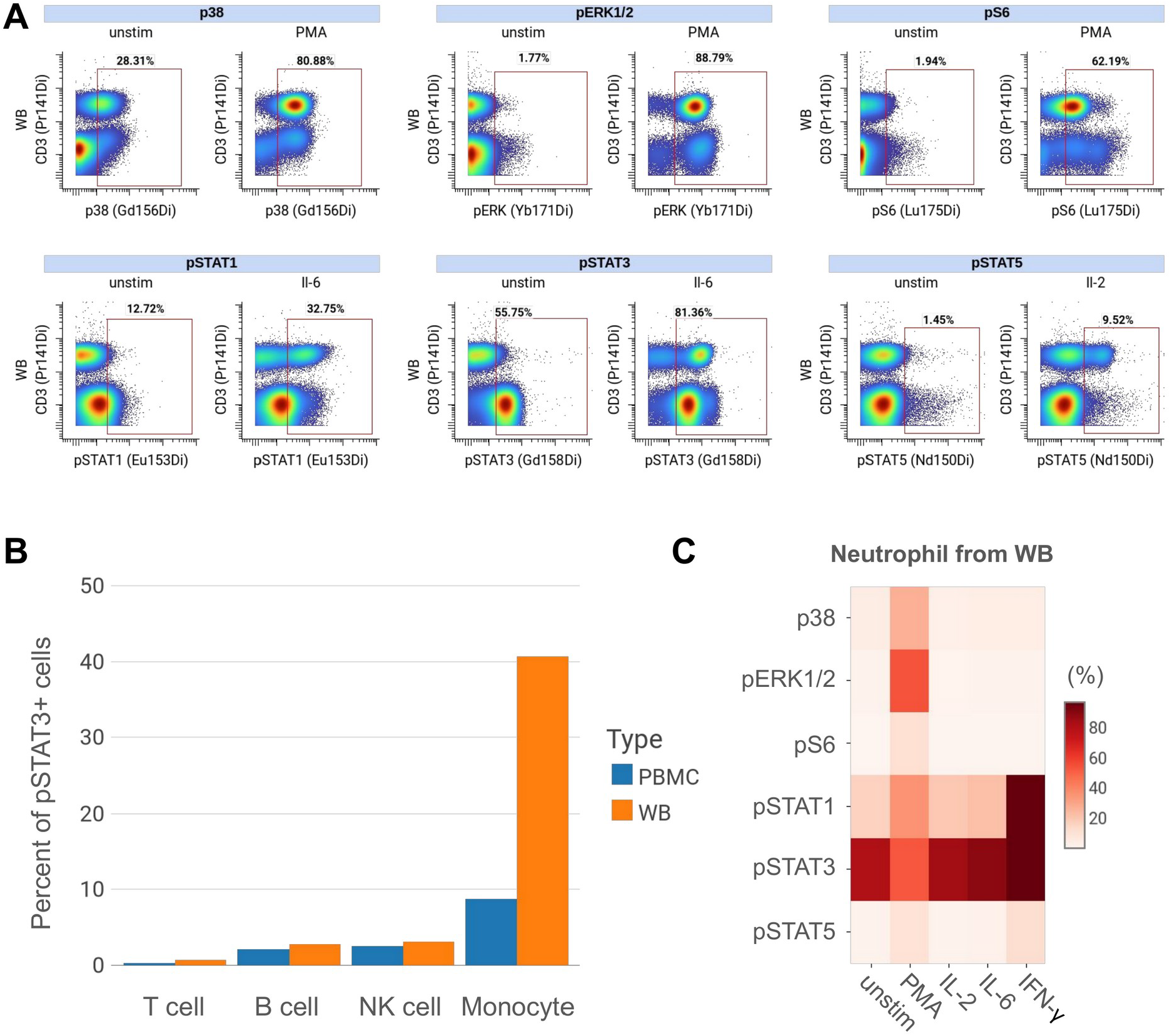
Basal and stimulation-induced phosphorylation in fixed whole blood and PBMCs. **(A)**Representative biaxial plots of p38, pERK1/2, pS6, pSTAT1, pSTAT3, and pSTAT5 induction, comparing unstimulated and stimulated conditions (PMA/ionomycin, IL-2, IL-6, or IFN-γ) in whole blood. Percentages indicate the fraction of cells positive for the indicated phosphorylation marker within the gated leukocyte population. **(B)** Representative bar plot illustrating frequencies of pSTAT3-positive events in major cell populations (T, B, NK and monocyte) in unstimulated PBMCs and whole blood from a single healthy donor. **(C)** Heatmap illustrating gated frequencies of phosphorylation markers in neutrophils from whole blood samples under unstimulated and stimulated conditions (PMA/ionomycin, IL-2, IL-6, or IFN-γ).

Responses differed by subset (Figure 3). PMA/ionomycin induced pS6 most strongly in B cells, and pERK1/2 in monocytes.IL-2 induced pSTAT5 in NK and T cells, while IL-6 elicited pSTAT3 in T cells and monocytes. IFN-γ increases pSTAT1 in monocytes, neutrophils, and B cells, along with pSTAT5 in monocytes. These activation profiles serve as internal controls for panel performance and reagent reactivity.

**Figure 3.**
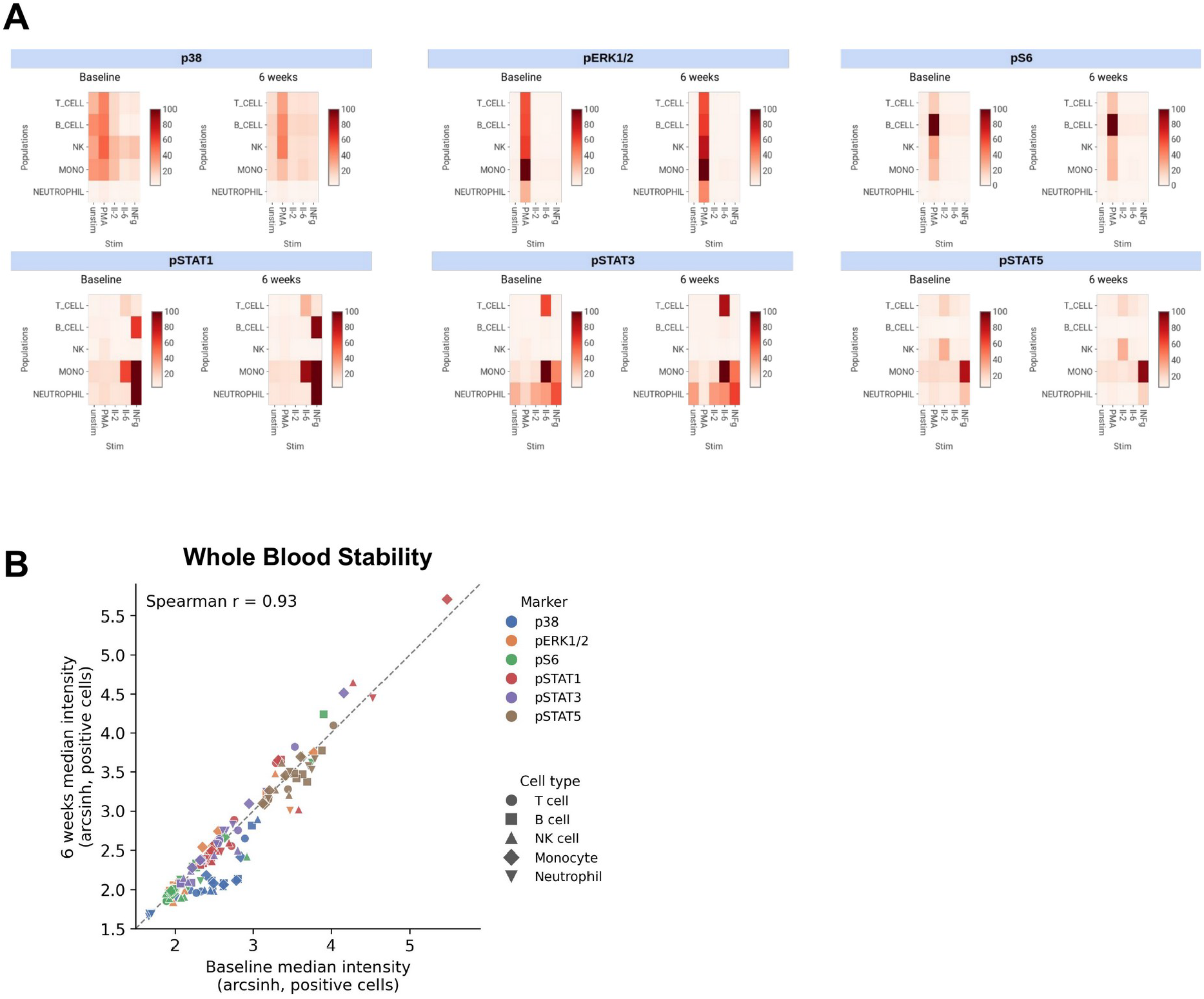
Storage stability of stimulation-induced signal. **(A)** Heatmaps show the percentage of cells positive for p38, pERK1/2, pS6, pSTAT1, pSTAT3, and pSTAT5 across five immune populations (T cells, B cells, NK cells, monocytes, and neutrophils) and five stimulation conditions (unstimulated, PMA/ionomycin, IL-2, IL-6, and IFN-γ) in whole blood at baseline and after 6 weeks of storage. **(B)** Median signal intensity of phosphoprotein-positive cells (arcsinh-transformed, cofactor 5) at 6 weeks versus baseline for all 150 marker, population, and stimulation combinations in (A). Colors indicate markers and shapes indicate cell types. Dashed line, identity.

### Fixed whole blood preserves neutrophil and monocyte phospho-signals lost in PBMCs

The difference between sample types was specific to monocytes. Basal pSTAT3-positive frequencies in T, B, and NK cells differed by less than 1 percent between whole blood and PBMCs. Monocytes differed by about 32 points, with ∼41% pSTAT3-positive cells in whole blood versus ∼9% in PBMCs (Figure 2B).

Neutrophils, which PBMC isolation removes, had ∼75–80% pSTAT3-positive cells without stimulation. IFN-γ raised pSTAT1-positive neutrophils from ∼25% to over 90%, and PMA/ionomycin raised pERK1/2-positive neutrophils to ∼60% (Figure 2C).

### Phospho-signals in fixed whole blood are retained after 6 weeks of storage

We measured the median channel value for each of the six phosphoproteins across five immune populations and five stimulation conditions in whole blood at baseline and after 6 weeks of storage (Figure 3A). Median channel values at 6 weeks correlated with baseline across all 150 combinations (Spearman r = 0.93; Figure 3B), and 119 of 150 were within 20% of baseline. Correlations were highest in neutrophils (r = 0.99) and T cells (r = 0.98) and ranged from 0.85 to 0.89 in B cells, NK cells, and monocytes. All of the subset-specific stimulation patterns described above were present at 6 weeks, and for all six markers the highest median channel value occurred in the same population and condition at both time points. p38 was the least stable marker, retaining 70% of its baseline median channel value; excluding p38, the correlation was r = 0.98. Stabilized whole blood can therefore be biobanked, shipped from remote clinical sites, and batch-processed centrally.

## Discussion

Fixing whole blood at the point of collection removes the PBMC isolation and freeze-thaw steps that alter phosphorylation before measurement. Conventional phospho-flow workflows isolate PBMCs by density-gradient centrifugation before fixation, adding hours of ambient handling and mechanical stress. In our data, this processing removed neutrophils and reduced the basal pSTAT3-positive monocyte fraction from ∼41% to ∼9%, while T, B, and NK cells were unchanged.

Because fresh PBMCs were not profiled, our design cannot separate the effects of density-gradient isolation from those of cryopreservation and thaw. Both steps are reported to alter phospho-signaling and subset recovery,^3,4^ and stabilized whole blood avoids both. Phospho-epitopes in fixed whole blood were stable for at least 6 weeks, a window long enough for batched acquisition and for shipment from remote sites, although shipping itself was not tested. Separately, TokuKit-fixed whole blood preserves immune population frequencies for up to 24 months at −80°C by mass cytometry (Pearson r = 0.998 versus baseline; n = 3 donors).^7^ Phospho-epitope stability beyond 6 weeks was not tested here. Because both sample types were profiled on the same panel, signals can be compared directly between stabilized whole blood and cryopreserved PBMC. Relative to fresh whole blood fixation ^5^, the protocol trades same-day staining for storage and batching, which removes the need for a staining laboratory at each collection site.

### Limitations

Methanol permeabilization, required to access intracellular phospho-epitopes, damages cell membranes and limits recovery to approximately 20% of input cells. This limits use with low-input samples, such as pediatric draws or small-volume research collections, where too few cells may remain to resolve rare populations after gating. The antibody panel was validated under this fixation and permeabilization workflow only. Because methanol destroys some antibody-binding epitopes, clones outside the validated panel and alternative phospho-targets require independent re-validation. Finally, the whole blood-versus-PBMC comparisons and the 6-week storage data come from a single donor and should be treated as proof of concept. Donor-to-donor variability in baseline signaling and in PBMC-processing artifacts has not been assessed, so replication in a larger cohort is needed to confirm the reproducibility and generalizability of these findings. We also did not include a fresh, immediately fixed reference sample or a head-to-head comparison with other whole blood stabilizers, so this study makes no equivalence claims against those methods.

## Resource availability

### Lead contact

Further information and requests for resources and reagents should be directed to and will be fulfilled by the lead contact, Li-Chun Cheng.

### Technical contact

Technical questions on executing this protocol should be directed to and will be answered by the technical contact, Li-Chun Cheng.

### Materials availability

TokuKit is commercially available from Teiko and through Fisher Scientific (NC3900747). The 20-plex palladium barcoding kit was prepared in-house following Rybakowska et al.^6^ Metal-conjugated antibodies prepared in-house (Table S2) were made with the commercial conjugation kits listed in the key resources table.

### Data and code availability

Data reported in this paper will be shared by the lead contact upon request. This paper does not report original code.

Any additional information required to reanalyze the data reported in this paper is available from the lead contact upon request.

## Acknowledgments

This work was funded by Teiko.

## Author contributions

L.S. performed the experiments, analyzed the data, prepared the figures, and wrote the original draft.

M.C. and C.L. optimized the protocol. L.-C.C. prepared the figures and reviewed and edited the manuscript. A.H. and J.J. reviewed and edited the manuscript.

## Declaration of interests

L.S., M.C., C.L., A.H., J.J., and L.-C.C. are employees and shareholders of Teiko. TokuKit is a commercial product developed and distributed by Teiko.

## Key resources table

| REAGENT or RESOURCE | SOURCE | IDENTIFIER |
|---|---|---|
| Antibodies |  |  |
| pSTAT1 [58D6]<sup>153</sup>Eu | Standard BioTools | Cat#201309 |
| pSTAT3 [4/P-STAT3]<sup>158</sup>Gd | Standard BioTools | Cat#201309 |
| pSTAT5 [47]<sup>150</sup>Nd | Standard BioTools | Cat#201309 |
| p38 [D3F9]<sup>156</sup>Gd | Standard BioTools | Cat#201309 |
| pERK1/2 [D13.14.4E]<sup>171</sup>Yb | Standard BioTools | Cat#201309 |
| pS6 [N7-548]<sup>175</sup>Lu | Standard BioTools | Cat#201309 |
| CD45 [HI30] | BioLegend | Cat#304002 |
| CD3 [UCHT1] | BioLegend | Cat#300438 |
| CD4 [SK3]<sup>174</sup>Yb | Standard BioTools | Cat#3174004C |
| CD8 [SK1] | BioLegend | Cat#344702 |
| CD20 [2H7] | BioLegend | Cat#302302 |
| CD56 [NCAM16.2] | BD | Cat#559043 |
| CD16 [B73.1] | BioLegend | Cat#360702 |
| CD14 [M5E2] | BioLegend | Cat#301862 |
| CD11c [Bu15]<sup>147</sup>Sm | Standard BioTools | Cat#3147008B |
| CD123 [6H6] | BioLegend | Cat#306002 |
| CD66b [G10F5] | BioLegend | Cat#305102 |
| CD235ab [HIR2] | BioLegend | Cat#306602 |
| Biological samples |  |  |
| Peripheral blood collected in sodium heparin tube | Healthy volunteer | N/A |
| Peripheral blood mononuclear cell (PBMC) prepared from healthy volunteer | Stanford Blood Center | BW019 |
| Chemicals, peptides, and recombinant proteins |  |  |
| Human TruStain FcX | BioLegend | Cat#422302 |
| RPMI 1640 medium with L-glutamine | Thermo Fisher Scientific | Cat#11875093 |
| Fetal bovine serum | Thermo Fisher Scientific | Cat#16140071 |
| Antibiotic-Antimycotic (100×) | Thermo Fisher Scientific | Cat#15240096 |
| Pierce Universal Nuclease for Cell Lysis, 250 KU/mL | Thermo Fisher Scientific | Cat#88700 |
| Cell Stimulation Cocktail (PMA/ionomycin, 500×) | Thermo Fisher Scientific (eBioscience) | Cat#00-4970-93 |
| Recombinant human IL-2 | Thermo Fisher Scientific | Cat#PHC0026 |
| Recombinant human IL-6 | BioLegend | Cat#570804 |
| Recombinant human IFN-γ | BioLegend | Cat#570202 |
| Paraformaldehyde, 16%, methanol-free | Electron Microscopy Sciences | Cat#15710 |
| Methanol, HPLC grade | Fisher Chemical | Cat#A452-1 |
| Trypan blue, 0.4% | Thermo Fisher Scientific | Cat#T10282 |
| 10× Maxpar Barcode Perm (saponin) | Standard BioTools | Cat#201057 |
| 10× eBioscience permeabilization buffer | Thermo Fisher Scientific | Cat#00-8333-56 |
| Iridium intercalator, 500 µM | Standard BioTools | Cat#201192B |
| Maxpar Cell Acquisition Solution Plus | Standard BioTools | Cat#201244 |
| EQ four-element calibration beads | Standard BioTools | Cat#201078 |
| Critical commercial assays |  |  |
| TokuKit whole blood preservation kit | Teiko; also distributed by Fisher Scientific | Cat#NC3900747 |
| 20-plex Pd barcoding kit | Prepared in-house following Rybakowska et al.<sup>6</sup> | N/A |
| X8 conjugation kit | Standard BioTools | Cat#201300 |
| MCP conjugation kit | Standard BioTools | Cat#201106A |
| Software and algorithms |  |  |
| CellEngine | CellCarta | https://cellengine.com |
| Other |  |  |
| Helios mass cytometer | Standard BioTools | N/A |
| BD Vacutainer sodium heparin blood collection tubes | Fisher Scientific | Cat#02-689-5 |
| Countess cell counting slides | Thermo Fisher Scientific | Cat#C10228 |
| Countess 3 | Invitrogen | Cat#A49862 |
| Centrifuge | Thermo Fisher Scientific | Cat#75-009-261 |
| Round-Bottom Polystyrene Test Tubes with Cell Strainer Snap Cap | Fisher Scientific | Cat#08-771-23 |
| 15 and 50 mL conical tubes | Fisher Scientific | Cat#H13576781, Cat#H13576773 |
| Cluster tubes | Corning | Cat#4401 |

### Experimental model and study participant details Institutional permissions

All human blood samples analyzed in this study were collected from consenting healthy adult donors under IRB protocol #20250604, approved by WCG Clinical Services.

### Method details

Detailed step-by-step method details, including materials and equipment and troubleshooting, can be found in Document S1.

The workflow has four stages: stimulation and fixation of whole blood or PBMC (∼1 h); palladium barcoding and pooling (∼2 h); surface staining, methanol permeabilization, and intracellular staining (∼4 h); and CyTOF acquisition and analysis (1–3 h).

### Quantification and statistical analysis

Mass cytometry data were normalized to EQ four-element calibration beads and debarcoded. Events were gated sequentially on Gaussian parameters and event length, then on singlets and leukocytes, before identification of major immune subsets (Figure S1). Data were analyzed in CellEngine (CellCarta). Signal was quantified as the median signal intensity per subset. Stimulation responses were computed as the arcsinh ratio or fold change relative to the unstimulated control. Storage stability was assessed by comparing baseline (T0) with 6-week samples.

### Supplemental information

**Document S1**. Step-by-step method details, materials and equipment, troubleshooting, Figure S1, and Tables S1 and S2

## Notes

### Competing Interest Statement

The authors have declared no competing interest.

