## Supplemental Information for "Onsite fixation preserves phosphoprotein profile in human whole blood"

#### Materials and equipment

##### Buffer Preparation

**Resting media**

| Component | Final concentration | Amount |
|---|---|---|
| RPMI 1640 with L-glutamine | N/A | 49 mL |
| Fetal bovine serum | 1% | 0.5 mL |
| Antibiotic-Antimycotic (100×) | 1% | 0.5 mL |
| Total | | 50 mL |

**Prepare fresh on the day of use and hold at 37°C.**

**Nuclease buffer**

| Component | Final concentration | Amount |
|---|---|---|
| Cell Staining Buffer | N/A | 1000 µL |
| $MgCl_2$, 1 M (1000×) | 1 mM | 1 µL |
| Pierce Universal Nuclease (10,000×) | 25 U/mL | 0.1 µL |
| Total | | ~1 mL |

**Prepare fresh and warm to 37°C before use. Prepare 900 µL per sample.**

**1× Maxpar Barcode Perm (saponin)**

| Component | Final concentration | Amount |
|---|---|---|
| 10× Maxpar Barcode Perm | 1× | 300 µL |
| 1× PBS | N/A | 2700 µL |
| Total | | 3 mL per sample |

**1× eBioscience permeabilization buffer**

| Component | Final concentration | Amount |
| --- | --- | --- |
| 10× eBioscience Perm buffer | 1× | 200 µL |
| Type 1 water | N/A | 1800 µL |
| Total | | 2 mL per sample |

**Prepare 10% excess to allow for pipetting loss.**

**Iridium intercalator solution**

| Component | Pooled <10 samples | Pooled 10–20 samples |
| --- | --- | --- |
| 4% PFA in 1× PBS | 1 mL | 1 mL |
| Iridium, 500 µM | 0.2 µL (1:5000) | 0.5 µL (1:2000) |

**Cell Staining Buffer (CSB)**

Prepare the following materials:

| Item | Order number |
| --- | --- |
| PES 0.2 µm filter | Fisherbrand FB12566510 |
| Disposable Polystyrene Sterile Bottles, 1 L | Corning™ 430518 |
| 10× PBS, pH 7.2 | Rockland Immunochemicals MB0084000 |
| 30% BSA | Sigma Aldrich A728450ML |
| Type 1 water | Prepared in-house |

| | Quantity for 1 L |
| --- | --- |
| 10× PBS | 100 mL |
| 30% BSA | 16.7 mL |

| | |
|---|---|
| Type 1 water | Fill to 1000 mL |

**PEB**

Prepare the following materials:

| Item | Order number |
|---|---|
| PES 0.2 µm filter | Fisherbrand FB12566510 |
| Disposable Polystyrene Sterile Bottles, 1 L | Corning™ 430518 |
| 10× PBS, pH 7.2 | Rockland Immunochemicals MB0084000 |
| 0.5M EDTA, pH 8.0 (100X) | Thermo Fisher 15575020 |
| 30% BSA | Sigma Aldrich A728450ML |
| Type 1 water | Prepared in-house |

| | Quantity for 1 L |
|---|---|
| 10× PBS | 100 mL |
| 0.5M EDTA (100X) | 10 mL |
| 30% BSA | 16.7 mL |
| Type 1 water | Fill to 1000 mL |

**For all buffers:**

1. Mix liquids in a graduated cylinder or bottle with a magnetic stir bar.
2. Filter the solution through a PES 0.2 µm filter into a clean bottle(s).
    a. **Note:** filtration here is to remove particles from the solution. Does not need to be handled inside the BSC.
3. Label the solution with solution name, operator initials, preparation date, expiration date (6 months), and "filtered" on the bottle.
4. Store the solution at 4°C.

### Step-by-step method details

#### PBMC Sample Preparation

**Timing: Day before experiment**

1. Transfer cryopreserved PBMCs from liquid nitrogen to an ice bucket filled with dry ice. Quickly untwist the cryovial lid half-turn to release pressure; once nitrogen has evaporated, screw the cap finger tight.
2. Thaw PBMCs quickly in a 37°C water bath for approximately 90 seconds:
    a. Visually check after the first minute.
    b. Invert the sample three times to loosen up the core.
    c. Return to the water bath until only a small piece of ice remains.
3. Spray 70% ethanol on the cryovial and transfer to the BSC.
4. Transfer all cells into a 15 mL conical tube using a P1000 pipette.
5. Add 1 mL of warm cRPMI to the empty cryovial to collect remaining cells; transfer drop-wise to the tube.
6. Slowly add 3 mL warm cRPMI; wait one minute. Then slowly add 5 mL more (total ~10 mL).
7. Centrifuge at 300 × g, 8 minutes, room temperature. Aspirate supernatant, leaving ~100 µL.
8. Resuspend in 1 mL warm cRPMI + nuclease. Pipette up and down 10 times.

Note: If sticky cell clumps are present, add 9 mL cRPMI + nuclease and incubate for 5 minutes. If clumping persists, pass through a 70 µm cell strainer.

9. Add 9 mL warm cRPMI + nuclease and repeat centrifugation (300 × g, 8 min, RT). Aspirate, leaving ~100 µL. Total: two cRPMI washes.
10. Resuspend cells in 1 mL warm resting media. Take 10 µL for cell counting (Countess with trypan blue).
11. Ideal counting density: 0.5–10 million cells/mL. Dilute and recount if outside range.
12. After cell counting, resuspend PBMCs in resting media (RPMI + 1% FBS + 1% AA) at 2 million cells/mL in a 50 mL Falcon tube or appropriate vessel.
13. Place cells in the 37°C / 5% $CO_2$ incubator overnight (12–16 hours).

#### Whole blood preparation

14. Collect blood in sodium heparin tubes. Record collection time. Use within 4 hours of collection; do not exceed 24 hours.
15. Gently invert the sodium heparin tube 8–10 times to remix cells before aliquoting.
16. Aliquot 250–500 µL of blood per stimulation condition into labeled tubes.

Note: WB does not require an overnight rest step. Proceed directly to stimulation.

#### Whole blood and PBMC stimulation

**Timing: 30 minutes per condition (see Table S1)**

17. PMA/ionomycin: Cell Stimulation Cocktail (1:500) Target markers: pERK1/2, pS6, p38 | Incubation: 30 min (WB and PBMCs)
    a. Thaw Cell Stimulation Cocktail (500×) to room temperature immediately before use.
    b. Spike in 1 µL of cocktail per 500 µL of cell suspension (1:500). Mix by pipetting.
    c. Incubate at 37°C / 5% $CO_2$ for 30 minutes. Record time in/out.
    d. Immediately proceed to fixation.
18. IL-6 (100 ng/mL working solution) Target marker: pSTAT3 | Incubation: 30 min (WB and PBMCs)
    a. Prepare 100 ng/mL working solution on ice by diluting IL-6 stock in resting media. Refer to lot COA for stock concentration.
    b. Spike working solution into the cell suspension to achieve 100 ng/mL final concentration. Mix gently.
    c. Incubate at 37°C / 5% $CO_2$ for 30 minutes. Record time in/out.
    d. Immediately proceed to fixation.
19. IL-2 (100 ng/mL working solution) Target marker: pSTAT5 | Incubation: 30 min (WB and PBMCs)
    a. Prepare 100 ng/mL working solution on ice: dilute stock 1:200 in resting media.
    b. Spike working solution into the cell suspension to achieve 100 ng/mL final. Mix gently.
    c. Incubate at 37°C / 5% $CO_2$ for 30 minutes. Record time in/out.
    d. Immediately proceed to fixation.
20. IFN-γ (100 ng/mL working solution, 1:1000 from 0.1 mg/mL stock) Target markers: pSTAT1 | Incubation: 30 min (WB and PBMCs)
    a. Prepare 100 ng/mL working solution on ice: dilute 0.1 mg/mL stock 1:1000 in resting media.
    b. Spike working solution into the cell suspension to achieve 100 ng/mL final. Mix gently.
    c. Incubate at 37°C / 5% $CO_2$ for 30 minutes. Record time in/out.
    d. Immediately proceed to fixation.

#### Fixation of PBMC and whole blood

**Timing: 30 minutes**

21. PBMC fixation
    a. Prepare filtered 16% PFA; keep in the dark and on ice.
    b. Chill the centrifuge to 4°C while waiting for the PFA incubation.
    c. Add 1/10 vol of 16% PFA (final concentration = 1.6%) directly to the cells in media (0.1 mL PFA per 0.9 mL of cells). Pipet up and down to mix well and incubate for 10 minutes at room temperature.
    d. Once fixation is complete, pellet cells by centrifugation at 500 × g, 5 min, 4°C.

e. Aspirate supernatant (contains fixative), leaving 100 µL behind. Vortex briefly to disperse the pellet.
f. Wash cells with PEB buffer (1 mL per 1–2 million cells). Pellet at 500 × g, 5 min, 4°C.
g. Aspirate supernatant, leaving 100 µL behind. Vortex briefly to disperse. Repeat once more for two PEB washes total.
h. Resuspend cells in 10% DMSO/CSB at 10 million cells/mL.
i. Transfer to a −80°C freezer.

Note: Consider aliquoting at least 2 million cells per cluster tube, especially if cells will undergo methanol perm. If viability is below 50%, aliquot 2–3 million total cells per tube.

22. Whole blood fixation (TokuKit)
    a. Add Buffer One at 1.4× volume of blood, invert 10×, and wait 15 minutes at room temperature.
    b. Add Buffer Two at 4× volume of blood and invert 10×.
    c. Transfer to a −80°C freezer.

#### Thaw, nuclease treatment, and cell number normalization

**Timing: 1 hour**

This step is for preparing samples prior to staining.

23. Remove fixed frozen samples from −80°C and thaw at room temperature for 10 min. Once completely thawed, place on ice.
    a. For PBMCs, prepare a nuclease buffer and warm it to 37°C. Add 900 µL to each sample with a P1000 multichannel pipette and mix by pipetting three times. For whole blood, wash with CSB 3 times with 1x volume of whole blood and pellet cells 500 × g for 5 min at 4°C.
    b. Incubate 5 min to reduce stickiness from debris.
    c. Mix again by pipetting three times, then remove 10 µL for counting. Keep the remainder on ice. Count with trypan blue on an automated counter for fewer than 20 samples, or by flow cytometer for larger numbers.
    d. Normalize each sample to 0.7–1.2 × $10^6$ cells.

CRITICAL: Consistent cell number per sample is required for consistent barcode intensity across the series. Where a sample exceeds 1.2 × $10^6$ cells, either aliquot 1 × $10^6$ into a fresh tube and re-freeze the remainder in 10% DMSO, or discard the excess. Where a sample falls below 0.7 × $10^6$, pool additional aliquots of the same sample from the same processing lot.

#### Sample barcoding

**Timing: 1 hour**

This step is for barcoding samples to pool.

24. Sample barcoding
    a. Thaw the barcode set at room temperature. Prepare 1× Maxpar Barcode Perm (saponin) and 1× eBioscience Perm buffer and keep on ice.
    b. Pellet cells at 500 × g for 5 min at 4°C and remove the supernatant. Vortex to disperse the pellet.
    c. Add 1 mL of 1× saponin buffer, pellet at 500 × g for 5 min at 4°C, and remove supernatant. Vortex to disperse. Repeat for two saponin washes total.
    d. Aliquot 400 µL of 1× saponin buffer per well of a 96-well deep-well plate, one well per barcode. Keep the plate on ice.

Note: For whole blood samples, aliquot 900 µL per well.

    e. Transfer 5 µL of barcode solution to each well with a P10 multichannel pipette and mix with the saponin buffer. For whole blood, transfer 10 µL per well.
    f. Resuspend cells thoroughly with a multichannel pipette to ensure no clumps remain, then transfer the entire barcode/saponin volume to the cells. Mix by pipetting up and down.

CRITICAL: Both barcode and cells must be fully resuspended before combining. Incomplete resuspension is the most common cause of poor barcode separation and cannot be corrected downstream.

    g. Incubate 30 min at room temperature on a horizontal shaker at 300 rpm. Record start and end times.

Note: Assign barcodes so that all stimulation conditions from a single donor occupy the same barcode series. This places every within-donor comparison inside a single staining reaction and removes staining batch effects from the comparison that matters most.

    h. Pellet at 500 × g for 5 min at 4°C and remove supernatant. Vortex briefly.
    i. Add 1 mL of 1× eBioscience Perm buffer, pellet at 500 × g for 5 min at 4°C, and remove supernatant. Vortex briefly.
    j. Add 1 mL of 1× eBioscience Perm buffer and incubate 10 min at room temperature on a horizontal shaker before pelleting at 500 × g for 5 min at 4°C. Remove supernatant and vortex briefly. Two eBioscience Perm washes total.
    k. Add 1 mL of CSB, pellet at 500 × g for 5 min at 4°C, and remove supernatant. Vortex briefly.
    l. Combine samples from each barcode series into one 15 mL tube labeled with date and project identifier. Transfer cells from cluster tubes using a single P200 tip for all samples in the series, to minimize loss from cells adhering to dry plastic.
    m. Add 1 mL of CSB to the first cluster tube, pipette to recover cells from the tube wall, and transfer to the second tube. Repeat sequentially through the series, then combine into the 15 mL tube. Pellet at 500 × g for 5 min at 4°C and remove supernatant down to the cell volume required for antibody staining.

Note: Mark the target cell volume on the tube with a marker before pelleting, or load an empty tube with the target volume as a visual reference.

#### Surface staining, methanol permeabilization, and phosphostaining

**Timing: 5 hours**

This step is for the sequential staining and permeabilization of phosphomarkers.

25. Surface staining
    a. Add 1 µL of FcX block per $10^6$ cells. Vortex briefly and incubate 10 min at room temperature on a horizontal shaker at 300 rpm.
    b. Prepare the surface antibody cocktail in CSB. Filter through a 0.1 µm spin column if the cocktail has been through a freeze-thaw cycle. Add to the cells and incubate 60 min at room temperature on a shaker at 300 rpm. Record start and end times.
    c. Dilute the staining reaction at least tenfold with CSB: 1 mL for a 50–100 µL reaction, or 5 mL for a pooled 500 µL reaction.
    d. Pellet at 500 × g for 5 min at 4°C and remove supernatant. Vortex to disperse.
    e. Add 1 mL of CSB, pellet at 500 × g for 5 min at 4°C, and remove supernatant. Vortex to disperse. Two CSB washes total.

CRITICAL: Prepare on the day of staining. Do not use stored dilute PFA.

26. Methanol permeabilization and phosphostaining
    a. Prepare fresh 4% PFA in PBS by mixing 250 µL of 16% PFA with 750 µL of 1× PBS.
    b. Add 1 mL of 4% PFA/PBS, mix by pipetting, and incubate 10 min at room temperature.
    c. Centrifuge at 500 × g for 5 min at 4°C, aspirate, and vortex briefly.
    d. Wash once with 1 mL of PBS. Pellet at 500 × g for 5 min at 4°C and aspirate, leaving as little residual volume as possible.
    e. Add 1 mL of ice-cold 100% methanol, mix by pipetting, and incubate 10 min on ice.

CRITICAL: Set the centrifuge to 700 × g before proceeding. Methanol-permeabilized cells are sticky and fragile. Do not vortex at any point after this step. Switch to 1200 µL pipette tips and pre-wet each new tip by pipetting 1 mL of CSB up and down before touching cells. Cell loss during and after methanol permeabilization is the single largest source of yield variability in this workflow.

    f. Pellet at 700 × g for 5 min at 4°C and aspirate.
    g. Add 1 mL of CSB and pipette to resuspend. Pellet at 500 × g for 5 min at 4°C and aspirate. Repeat for two CSB washes total.
    h. Remove CSB down to the cell volume required for staining.
    i. Add the phospho/intracellular antibody cocktail. Incubate overnight at 4°C.
    j. Dilute the reaction at least tenfold with CSB, pellet at 500 × g for 5 min at 4°C, and remove supernatant. Vortex to disperse.
    k. Add 1 mL of CSB, pellet at 500 × g for 5 min at 4°C, and remove supernatant. Two CSB washes total.
    l. Add 1 mL of iridium intercalator solution and incubate 20 min at room temperature, or overnight at 4°C. Record start and end times.
    m. Pellet at 500 × g for 5 min at 4°C and remove supernatant.

CRITICAL: Do not freeze methanol-permeabilized samples after staining. Acquire on the same day.

#### Acquisition on mass cytometer

**Timing: approximately 1 h per 1–2 × $10^6$ cells**

This step is for the acquisition of stained samples on the mass cytometer.

27. Acquisition sample preparation
    a. Wash once with 1 mL of CSB.
    b. Prepare a fresh nuclease buffer and warm to 37°C. Add 1 mL and incubate 3 min at room temperature on a shaker.
    c. Centrifuge at 500 × g for 5 min and aspirate as completely as possible.
    d. Before washing, use a P200 pipette to remove as much residual CSB as possible. Then wash once with 2 mL of CAS-Plus and count cells.

CRITICAL: Wash methanol-permeabilized samples only once with CAS-Plus. Additional washes cause cell loss without benefit. Removing residual CSB before the wash prevents salt carryover into the instrument; salt accumulates on the cone, reducing sensitivity and increasing CV across the run.

    e. Resuspend in CAS-Plus supplemented with EQ beads at 1:50 of final volume, to a density of 0.7 × $10^6$ cells/mL. Recount if needed for accurate density.
    f. Filter twice through 5 µm cell strainers into a final FACS tube. Keep on ice until loading.
    g. Acquire on the mass cytometer.

### Troubleshooting

#### Problem 1 (PBMC thaw)

Low cell recovery immediately after thawing cryopreserved PBMCs

**Potential solution:** Confirm cells were thawed quickly (37°C water bath, ~90 sec) and diluted slowly with warm cRPMI to avoid osmotic shock. If sticky clumps are visible, add the full 9 mL cRPMI + nuclease wash and incubate 5 min before the second centrifugation rather than proceeding directly; pass through a 70 µm strainer if clumping persists.

#### Problem 2 (PBMC/whole blood fixation)

Cell loss during PFA fixation and PEB washes.

**Potential solution:** Ensure PFA is freshly prepared and filtered, and that centrifugation is performed at the specified 500 × g, 4°C for the full 5 min. Vortex briefly to fully disperse the pellet before each wash. Avoid aspirating below the 100 µL residual volume, as over-aspiration at this stage is a common source of pellet disturbance and loss.

#### Problem 3 (methanol permeabilization)

Low or inconsistent cell recovery after methanol permeabilization. This is the single largest source of yield variability in this workflow.

**Potential solution:** Increase the post-methanol centrifugation speed and duration if pellets are not fully recovered at 700 × g, 5 min. Switch to wide-bore or 1200 µL tips for all steps after methanol addition, pre-wet each new tip with 1 mL of CSB before it touches cells, and never vortex after methanol addition, as permeabilized cells are fragile and sticky. Minimize the total number of post-methanol washes to only those specified.

#### Problem 4 (barcode pooling)

Cell loss while combining cluster tubes into a single barcode series.

**Potential solution:** Use a single P200 tip for all samples in a series rather than switching tips, and recover cells adhering to the tube wall by rinsing sequentially with CSB through the series before combining, as specified.

#### Problem 5 (related to acquisition)

Instrument clogging or reduced sensitivity during acquisition due to sticky, methanol-permeabilized cells.

**Potential solution:** Methanol-permeabilized cells remain sticky even after rehydration and are prone to aggregating, which can clog the injector or foul the cone. Wash methanol-permeabilized samples only once with CAS-Plus, as specified. Use a P200 pipette to remove residual CSB as completely as possible before this wash, since carried-over salt compounds cone fouling. Filter the final resuspension twice through 5 µm cell strainers immediately before loading, and do not let the prepared sample sit for an extended period before acquisition, as clumping increases with time.

### Supplemental figure and tables

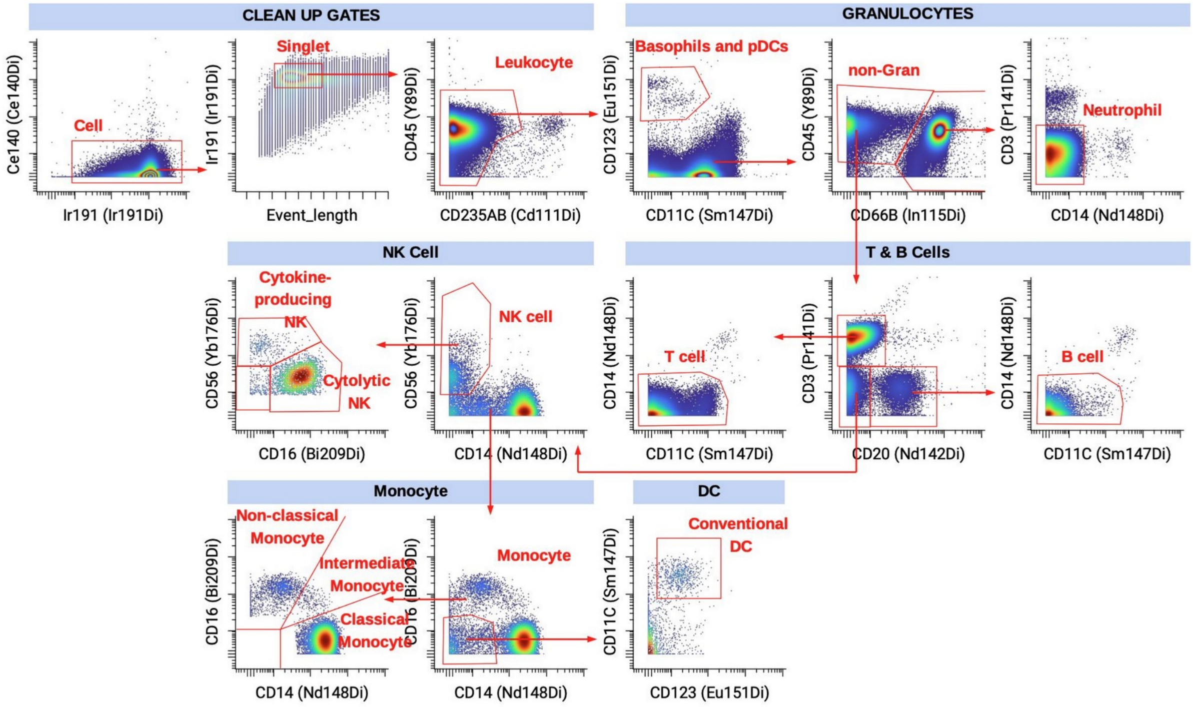


**Figure S1. Gating scheme, related to Figures 2 and 3 and STAR Methods.** Representative gating strategy for major immune subsets by mass cytometry. Clean-up gates (cell, singlet, leukocyte) are followed by gating of basophils and pDCs and of neutrophils using CD123, CD11c, CD45, and CD66b, and gating of non-granulocyte leukocytes into T cells, B cells, cytokine-producing and cytolytic NK cells, conventional DCs, and classical, intermediate, and non-classical monocytes using CD3, CD20, CD56, CD11c, CD123, CD16, and CD14.

**Table S1. Stimulation conditions**

| Condition | Stimulus | Working conc. | Dilution | Target markers | Incubation (WB / PBMC) |
|---|---|---|---|---|---|
| T0 | None (stabilized at collection) | N/A | N/A | Basal | 0 min |
| PMA + ionomycin | Cell Stimulation Cocktail (500×) | 1× | 1:500 | pERK1/2, pS6, p38 | 30 min / 30 min |
| IL-2 | Recombinant human IL-2 | 100 ng/mL | per lot | pSTAT5 | 30 min / 30 min |

| | | | | | |
|---|---|---|---|---|---|
| IL-6 | Recombinant human IL-6 | 100 ng/mL | per lot | pSTAT3 | 30 min / 30 min |
| IFN-γ | Recombinant human IFN-γ | 100 ng/mL | ~1:1000 | pSTAT1 | 30 min / 30 min |

**Table S2. Antibody panel**

| Target | Clone | Isotope | Company | Marker Purpose |
|---|---|---|---|---|
| pSTAT1 | 58D6 | $^{153}$Eu | Standard Biotools | Phosphomarker |
| pSTAT3 | 4/P-STAT3 | $^{158}$Gd | Standard Biotools | Phosphomarker |
| pSTAT5 | 47 | $^{150}$Nd | Standard Biotools | Phosphomarker |
| p38 | D3F9 | $^{156}$Gd | Standard Biotools | Phosphomarker |
| pERK1/2 | D13.14.4E | $^{171}$Yb | Standard Biotools | Phosphomarker |
| pS6 | N7-548 | $^{175}$Lu | Standard Biotools | Phosphomarker |
| CD45 | HI30 | $^{89}$Y | Prepared In-house | Total leukocyte |
| CD3 | UCHT1 | $^{141}$Pr | Prepared In-house | All T cells |
| CD4 | SK3 | $^{174}$Yb | Standard Biotools | Helper T cells |
| CD8 | SK1 | $^{112}$Cd | Prepared In-house | Cytotoxic T cells |
| CD20 | 2H7 | $^{142}$Nd | Prepared In-house | B cells |
| CD56 | NCAM16.2 | $^{176}$Yb | Prepared In-house | Natural Killer (NK) cells |
| CD16 | B73.1 | $^{209}$Bi | Prepared In-house | NK cells, neutrophils, and macrophages |
| CD14 | M5E2 | $^{148}$Nd | Prepared In-house | Monocytes and macrophages |
| CD11c | Bu15 | $^{147}$Sm | Standard Biotools | Dendritic cells |
| CD123 | 6H6 | $^{151}$Eu | Prepared In-house | Plasmacytoid |

| | | | | |
|---|---|---|---|---|
| | | | | dendritic cells and basophils |
| CD66b | G10F5 | $^{115}In$ | Prepared In-house | Neutrophils |
| CD235ab | HIR2 | $^{111}Cd$ | Prepared In-house | Red Blood Cells |
